# Inferring the Joint Demographic History of Multiple Populations: Beyond the Diffusion Approximation

**DOI:** 10.1101/103275

**Authors:** Julien Jouganous, Will Long, Simon Gravel

## Abstract

Understanding variation in allele frequencies across populations is a central goal of population genetics. Classical models for the distribution of allele frequencies, using forward simulation, coalescent theory, or the diffusion approximation, have been applied extensively for demographic inference, medical study design, and evolutionary studies. Here we propose a tractable model of ordinary differential equations for the evolution of allele frequencies that is closely related to the diffusion approximation but avoids many of its limitations and approximations. We show that the approach is typically faster, more numerically stable, and more easily generalizable than the state-of-the-art software implementation of the diffusion approximation. We present a number of applications to human sequence data, including demographic inference with a five-population joint frequency spectrum and a discussion of the transferability of demographic histories across populations.

## 1 Introduction

Understanding the role of demography and selection in shaping genetic diversity is a central challenge in population genetics. In recent years, genome sequencing experiments have generated large amounts of data that can be used to test and refine this understanding. Detailed models of human genetic diversity have shed a new light on the origins and history of modern humans, and have helped researchers design and interpret biomedical studies.

Present-day genomes depend on a large number of more or less randomly occurring mutations, recombinations, matings, and deaths. Individual events and their genomic consequences bear limited information about the past. We can, however, learn a lot by statistically integrating information across entire genomes and populations. Classical work has focused on simple summaries of genomic diversity that could be computed analytically, such as the number of pairwise differences between individuals, the number of segregating sites in a sample, or linkage disequilibrium [4]. In recent years, the ability to simulate genomes and the availability of data has led to a wide array of summaries designed to identify different evolutionary forces [22, 24, 23]. Here we consider the allele frequency spectrum (AFS): We compute the frequency of the derived allele at each site, and build a histogram of the number of sites observed at each frequency.

The AFS is a classical measure of diversity [16] that has seen a surge in popularity recently because of improved computational approaches to estimate it. Back-in-time approaches based on coalescence theory [7, 6, 14] can be extremely efficient for neutral simulations and can handle large number of populations, but they often become cumbersome or intractable in models with selection. Forward-in-time approaches tend to be more easily generalized to account for selection. Despite recent progress in discrete whole-population methods [13], most current approaches are based on the diffusion approximation initiated by Fisher and Wright in the early 1930’s [8, 31, 4, 16]. In this approach, the evolution of the AFS is described as a continuous process through a partial differential equation. Several simulating tools based on this diffusion approximation have been implemented and distributed to the population genetics community [12, 20, 21], among which *∂a∂i* is probably the most commonly used. In the past few years, these numerical tools have been used widely and led to significant results from both historical and biological points of view [12, 9, 25].

Despite these successes, computational cost and stability remains an important limitation. For instance, available software can only handle up to 4 populations simultaneously (*∂a∂i* can handle 3). The time needed to perform individual simulations can be prohibitive when many simulations must be run, for example when performing bootstrap analysis for multiple models. Furthermore, biases inherent to Kingman’s coalescent or the diffusion approximation, which were benign in small samples, can become important in contemporary datasets [1].

Here, we describe a new and efficient method to simulate the AFS that is more efficient and more numerically stable than the state-of-the-art, yet does not require the diffusion approximation. We integrate this simulation engine into the *∂a∂i* inference framework to facilitate inference from allele frequency distributions across a broader range of problems than was previously possible.

## 2 Method and material

### 2.1 Heterozygosity rate evolution

First consider a large population of *N* diploid individuals evolving under the neutral Wright-Fisher model. Generations are discrete and individuals from generation *k* + 1 receive alleles drawn randomly and with replacement from the parental alleles present in generation *k*. Because alleles in a diploid individual are inherited and transmitted independently in the neutral case, we can forget about diploid individuals and think of the population as a set of 2*N* haploid samples (or *ploids*, for short). We are interested in the expected number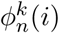 of sites where the alternate allele is observed exactly *i* times in a sample of size *n* at generation *k*. We neglect correlations between sites and suppose that each locus is transmitted independently.

In this model, heterozygosity is proportional to 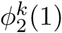, the number of heterozygous sites in sample of n=2 ploids. Under neutral Wright-Fisher reproduction, this follows the classical recursion

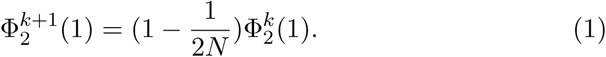

The derivation of this recursion is simple: If two ploids at generation *k* + 1 are inherited from the same parental ploid, which happens with probability 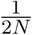, they are identical and do not contribute to 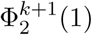. Otherwise (with probability 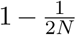) they are drawn randomly without replacement from the previous generation. The probability that they are different is 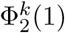, by definition, leading to Equation (1).

### 2.2 Neutral case AFS

We can similarly derive a recursion equation for the neutral AFS for arbitrary *n* and *i*. This time we draw *n* ploids from the previous generation. The probability that a given pair of ploids is still 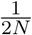. For simplicity, we suppose that at most one ploid pair shares a parent at each generation. The situation with multiple coalescent events per generation, which can occur for large sample sizes [1], is a straightforward generalization discussed in Section B.2.

If there is no coalescence, the *n* ploids at generation *k* + 1 are copied from *n* randomly drawn ploids at generation *k* and the distribution of allele frequencies does not change: 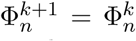. If there is a coalescence, the *n* descendants are copied from a random set of *n* − 1 parental haplotypes. The distribution of possible parental frequencies is described by 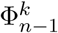. To treat all possible coalescence cases in a unified way, we imagine that we always draw *n* parental ploids, with the understanding that one selected parental ploid may not leave descendants. By doing this, we can express 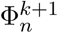 as a simple linear function of 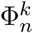:

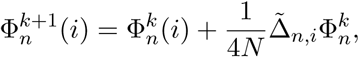
where 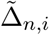 is a sparse linear operator describing the change in allele frequency due of drift in a single generation. Its coefficients are provided in Supplement A. It is related to other transition matrices derived in [5, 26] (more on these relationships in the discussion).

*De novo* mutations can change the allele type and therefore also affect the allele frequency. Their effects on the *i^th^* entry of the AFS is described by the linear operator *M_i_*:

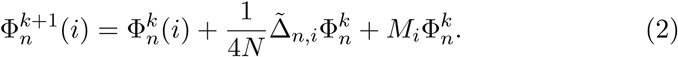

As a first approximation and to easily compare to commonly used simulation tools, we consider the infinite-sites model [17], in which the only mutations allowed are at previously invariant sites. Assuming that at most one mutation per generation per site occurs in the sample, we simplify the mutation term and write it as an additive source term: 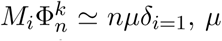 being the mutation rate. In the infinite-sites model, we therefore have

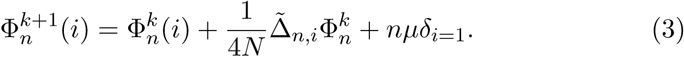

A more general form of the mutation operator, accounting for backwards mutations, is discussed in Appendix B.

Equation (3) is formally similar to a numerical discretization of the diffusion approximation. However, it avoids two approximations: first, the approximation of a discrete Wright-Fisher system by a continuous diffusion and, second, the approximation of a continuous diffusion by a discrete numerical partial differential equation (PDE) solver. Even though we use discrete frequency space, we will use a continuous time approximation that is easily simulated using classical integration schemes:

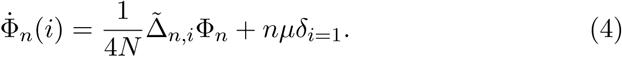

### 2.3 Modeling selection and dominance

To obtain a similar recursion equations under selection, we consider a model of selection in which ploids are drawn uniformly from the previous generation, but are accepted with different rates depending on the fitness of the parent. Alleles drawn from individual I are accepted with probability 1 for genotype *aa*, 1 + *hs* for genotype *aA*, and 1 + *s* for genotype *AA*: *s* is the selection coefficient, and *h* is the dominance. If a ploid is not accepted, it must be redrawn at random from the entire population.

In this model, contrary to the neutral case, we may need to draw more than *n* alleles from generation *k* to build 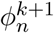. In what follows, we assume that *ns* ≪ 1, so that at most one allele is rejected from the sample at each generation. This assumption is also implicit in the diffusion approximation, but here again higher-order corrections can be computed.

Assuming *ns* ≪ 1, the recursion is (5):

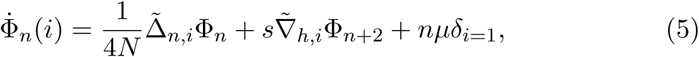

Where 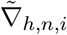 is a sparse linear operator describing the change in sample allele frequencies due to selection. It’s coefficients are given in Supplement A.

To compute 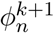, Equation (5) requires 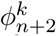, which itself requires 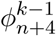, and so forth. The evolution equation for 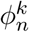 is not closed and cannot be solved directly in that form.

### 2.4 Moment closure method

We would like to find a closed approximation to Equation (5) that would make it possible to compute 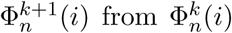. To do this, we want to approximate the larger samples AFS 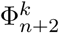 in terms of 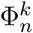. Intuitively, this should be possible for *n* large enough: We don’t learn very much more from a sample of size 102 than we do from a sample of size 100. In fact, the number of variants to be discovered in a sample of size 20*n* can be estimated accurately from a sample of size *n* by simple jackknife extrapolation [10]. The same kind of extrapolation - based on a jackknife method - can be successfully adapted to the much easier moment closure problem.

In this work, we considered a 3rd order jackknife to express entries of the higher order spectra Φ_*n*+1_ and Φ_*n*+2_ as linear combinations of 3 entries of Φ_*n*_. Higher order jackknives would increase accuracy at the cost of computational complexity and stability. Since jackknives provide a linear approximation, we can write 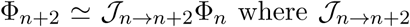 is a linear operator whose coefficients are provided in Appendix D. Under this approximation, the selection term becomes 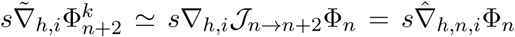, where we defined 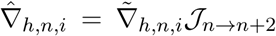 as the closure version of the selection operator. The resulting system is a closed and linear system of ordinary linear differential equations which can be solved using standard methods:

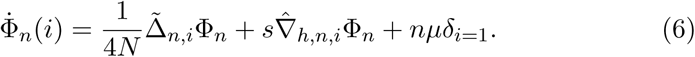

### 2.5 Multiple populations and migrations

In practice, we often want to study the distribution of allele frequencies across multiple populations where mating is more common within than across populations. In such a case, we consider the multidimensional AFS Φ_n_(**i**) where **n** is a vector of sample sizes of the different populations. Thus, Φ_*n*_1_,…, *n*_*p*__ (*i*_1_,…, *i*_*p*_) is the number of variants that are found *i*_1_ times in population 1, *i*_2_ times in population 2, and so forth.

Using the same method as for the single population case, we can derive a system of ordinary equations for the joint AFS Φ_n_ (see C.2). Because of migration, however, the evolution of Φ_n_ depends on Φ_ñ_ for **ñ ≠ n**: When creating *n_i_* ploids in population *i* and *n_j_* in population *j*, we may occasionally draw *n_i_* +1 ploids from *i* and *n_j_* − 1 from *j* (the case of strong migration, where we are likely the probability of drawing multiple migrants per generation, is discussed below). Thus evolution equations for Φ_n′_ is coupled for all **n′** such that *n′_j_* + *n′_k_* = *n_j_* + *n_k_*. Under weak migration, we can use a jackknife to write uncoupled approximations to these equations. A simple way of achieving this is to write the evolution equation for Φ_*n*_ in terms of the slightly larger AFS Φ_ń_ with **ń**_*j*_ = **n** + **e**_*j*_ and **e**_*j*_ a unit vector with a unit value for the *j*th population. We then have

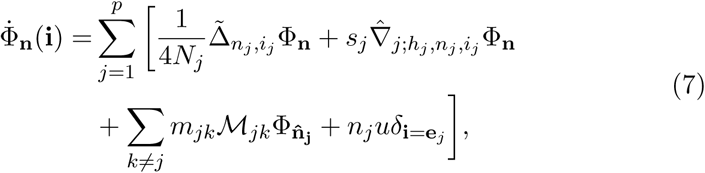
where 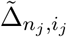 is the drift operator and 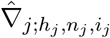 is the selection operator for population *j*, *ℳ_jk_* is the linear migration operator from populations *j* to population *k*, and the migration rate *m_jk_* is the proportion of ploids in population *k* whose parent was in population *j*. Parameters for the migration operator *ℳ_jk_* are provided in Appendix A.

We can then use a jackknife approximation as above to write

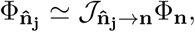

Where the sparse linear operator 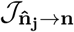 is a jackknife operator (The code distributed with this article uses a uses a slightly more complex jackknife formulation described in Appendix D). We can obtain a closed version of the migration operator, 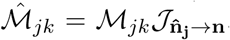, leading to a closed-form evolution equation for Φ_*n*_:

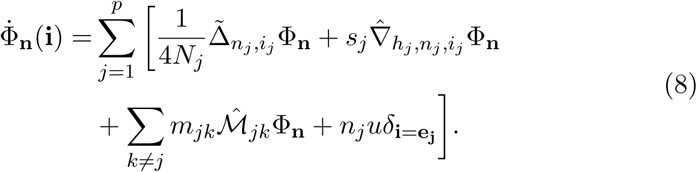

So far, we supposed that migration was weak enough that the expected number of migrants per generation in our sample was less than 1. However, many populations have received many migrants over few generations. This is the case, for example, in many populations from the Americas.

In such cases, we compute the resulting AFS directly without using the jackknife approximation. Because migration is a stochastic process, the number of migrant lineages varies across loci. A given sample of 10 ploids may derive 100% ancestry from population 1 at one locus, and 100% from population 2 at another. In the independent-sites model and two-way admixture, the distribution of lineages that trace back to one population is binomial: The expected frequency spectrum is an integral over the possible numbers of migrating lineages. To generate *n* admixed lineages, we can therefore need as many as n lineages from each of the source populations. We explain in Section B.3 how the number of source population lineages can be chosen to maximize computational efficiency.

### 2.6 Connections with the diffusion approximation

One of the standard approaches to simulate AFS evolution relies on a continuous approximation to the Wright-Fisher process. This approximation leads to an advection-diffusion equation describing the evolution of the allele frequencies density *ϕ*(*x*, *t*) (i.e. the expected proportion of alternate ploids at frequency *x* ∈ [0. 1] in the population and at time *t*).

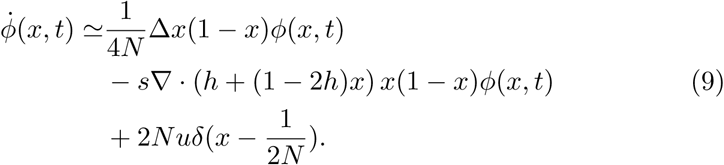

The most widely used tool based on this diffusion approximation is *∂a∂i* which numerically approximates the distribution *ϕ* with finite differences. The AFS entries are then computed via the integral:

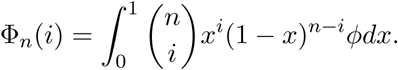

The AFS entries can be seen as moments of the distribution *ϕ* computed in a non-canonical basis of polynomials. The idea of deriving evolution equations for the classical moments 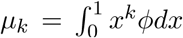 was proposed by Evans, Shvets and Slatkin [5]. However, the computation of the AFS from the moments *μ_k_* raises numerical instabilities as the formula involves alternating sums. A closed system of ordinary differential equations directly for the neutral Φ_*n*_(*i*) is evoked by the authors but deemed too complex to be of practical importance. The case with selection, lacking even a closed expression, is therefore even more daunting.

In fact, we show in Appendix C that by computing a moment equation for the diffusion equation, we obtain precisely Equation (5), which is not more complex than the original diffusion equation and is, as we will see, more numerically stable. The lack of closure of the evolution equation when selection is present is easily addressed by a jackknife procedure for the weak selective values consistent with the diffusion approximation.

### 2.7 Implementation and performance

We developed a python library, called *Moments*, to simulate multidimensional AFS and infer demographic history using the method described above. From the user perspective, the library is very similar to *∂a∂i*, since we reused the *∂a∂i* software architecture, including the user interface and data handling methods. We added a number of new convenience features, described in Appendix F, but the most important difference is performance.

Whereas *∂a∂i* could model up to three populations and *Multipop* [20] up to four, *Moments* can handle models with up to 5 populations with selection, migrations and population splits. The run-time is still exponential in the number of populations, so the direct computation of large sample sizes for more than three populations remains challenging.

To provide a fair comparison between *Moments* and *∂a∂i* for a comparable problem size, we need to consider both accuracy and computation time. *∂a∂i* users can chose the number of grid points used for solving the diffusion equation. A large number of points results in slower but more accurate integration. We used the recommended strategy in *∂a∂i*, namely using three different grid sizes and performing Richardson extrapolation

*Moments* also presents a tradeoff between speed and accuracy because the moment closure approximation improves with increasing *n*. *Moments* can therefore be made more accurate by computing the AFS for *n*′ larger than the desired *n*, and subsampling to *n* after the integration. For the simulations below, we simply integrated with the desired *n*.

Table 1 shows comparisons between the two methods on a few test cases. A more extensive set of test cases ranging from the analytically solvable equilibrium case to the numerically exacting is provided in the supplementary material E.

**Table 1:**
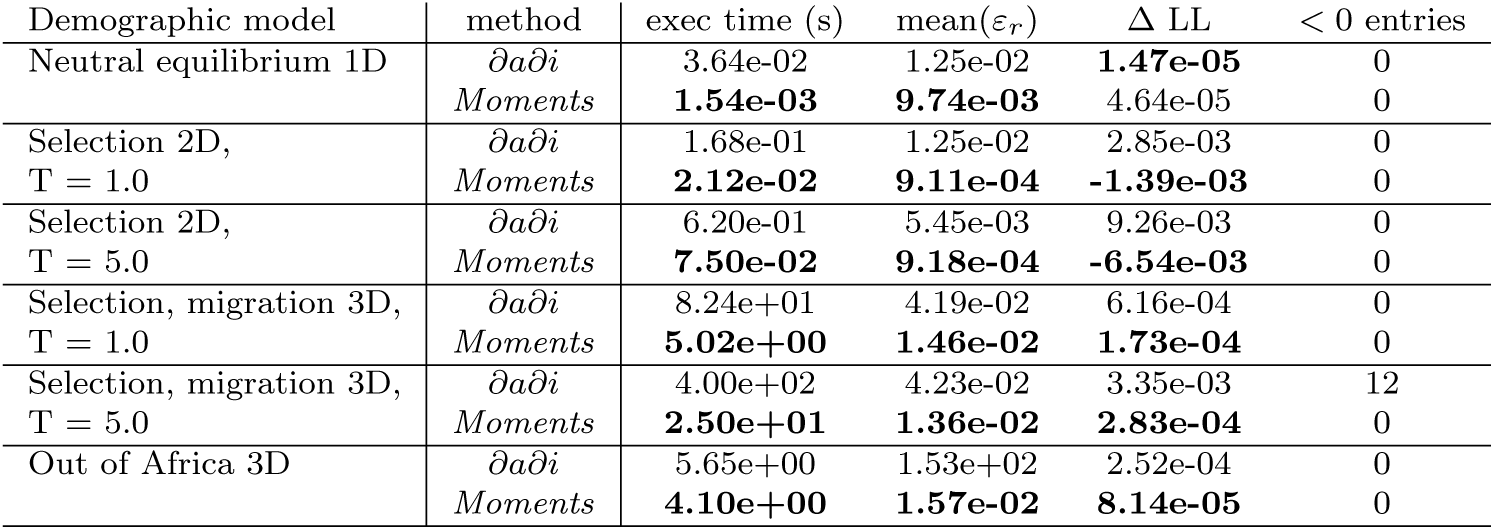
Performance comparisons between *∂a∂i* and *Moments* on several scenarios (30 samples per population). The full bench is provided in Appendix E. For *∂a∂i* simulations we used Richardson extrapolation to improve convergence. The time T provided is the simulation time in genetic units.

| Demographic model | method | exec time (s) | mean($\varepsilon_r$) | $\Delta LL$ | < 0 entries |
| --- | --- | --- | --- | --- | --- |
| Neutral equilibrium 1D | $\partial a \partial i$ | 3.64e-02 | 1.25e-02 | **1.47e-05** | 0 |
|  | *Moments* | **1.54e-03** | **9.74e-03** | 4.64e-05 | 0 |
| Selection 2D,<br>T = 1.0 | $\partial a \partial i$ | 1.68e-01 | 1.25e-02 | 2.85e-03 | 0 |
|  | *Moments* | **2.12e-02** | **9.11e-04** | **-1.39e-03** | 0 |
| Selection 2D,<br>T = 5.0 | $\partial a \partial i$ | 6.20e-01 | 5.45e-03 | 9.26e-03 | 0 |
|  | *Moments* | **7.50e-02** | **9.18e-04** | **-6.54e-03** | 0 |
| Selection, migration 3D,<br>T = 1.0 | $\partial a \partial i$ | 8.24e+01 | 4.19e-02 | 6.16e-04 | 0 |
|  | *Moments* | **5.02e+00** | **1.46e-02** | **1.73e-04** | 0 |
| Selection, migration 3D,<br>T = 5.0 | $\partial a \partial i$ | 4.00e+02 | 4.23e-02 | 3.35e-03 | 12 |
|  | *Moments* | **2.50e+01** | **1.36e-02** | **2.83e-04** | 0 |
| Out of Africa 3D | $\partial a \partial i$ | 5.65e+00 | 1.53e+02 | 2.52e-04 | 0 |
|  | *Moments* | **4.10e+00** | **1.57e-02** | **8.14e-05** | 0 |

In the case of neutral equilibrium with constant population size, we can compare our computations to the exact solution. For more complex cases, where analytical results are unavailable, we use *∂a∂i* with a very fine frequency grid and use the spectrum thus obtained as a reference to compare our method and *∂a∂i*. Generating the ‘truth’ set with *∂a∂i* can induce a bias in favor of *∂a∂i*. We consider several configurations with up to 3 dimensions including selection, migration and non constant populations sizes. We used 30 samples per population.

The first 5 cases are simple integrations starting from a null spectrum: a single population without selection until the equilibrium is reached, two isolated populations under selection and three populations with selection and migrations. The sixth model is the demographic model of Out of Africa expansion described in [9]. Details of the models, and additional benchmarks, are provided in Appendix E.

We used different metrics to compare the simulations such as the execution time and the mean relative error *ε_r_* compared to the ‘true spectrum’. We also considered the difference in likelihood Δ*LL* = log(*L*(*truth*, *truth*)) − log(*L*(*truth*, *test*)), where *L*(*x*, *y*) is the probability to observe the AFS *x* assuming that the expected AFS is *y*, ‘truth’ is the AFS computed with third order Richardson extrapolations using fine grids – 30, 35 and 40 times the sample size for 1D and 2D cases and 5, 6 and 8 times the sample size for 3D simulations – in *∂a∂i*, and ‘test’ is the AFS computed using each software with regular settings.

Finally, an issue with *∂a∂i* in large sample sizes is that the AFS occasionally returns impossible negative allele frequencies. For each test case we therefore also counted the number of negative AFS entries.

On these several examples, *Moments* performs better than *∂a∂i* for most metrics. In the more general set of benchmarks provided in Appendix E, we find a few instances where *∂a∂i* outperforms *moments*. Overall, we find that *Moments* is particularly efficient for integrations over long time periods, and is generally much more robust to negative entries.

For 4 and 5 populations models, we could not compare *Moments* to *∂a∂i*, since *∂a∂i* does not allow for such high-dimensional simulations.

## Application to data

### 3.1 3 populations Out of Africa

We used our method to update the Out of Africa expansion model described in [9, 12]. These models are used in a wide variety of medical and evolutionary applications.

We used autosomal synonymous sequence data from the publicly available Thousand Genome Project (1000G) [3, 27]. We first computed the joint AFS for three populations: Yoruba individuals in Ibadan, Nigeria (YRI); Utah residents with northern and western European ancestry (CEU); and Han Chinese from Beijin (CHB). We restricted our analysis to 80 ploids for each population and fit the same 13-parameter demographic model as in [12, 9]. However, whereas previous studies had access to capture data for a subset of the exome, here we had access to the entire high-coverage exome data. Moreover, we updated the mutation coefficient and the generation time to more realistic values: respectively *μ* = 1.44 × 10^−8^ [11] and *T_g_* = 29*y* [28]. The best-fit model is represented in Figure 1.

**Figure 1:**
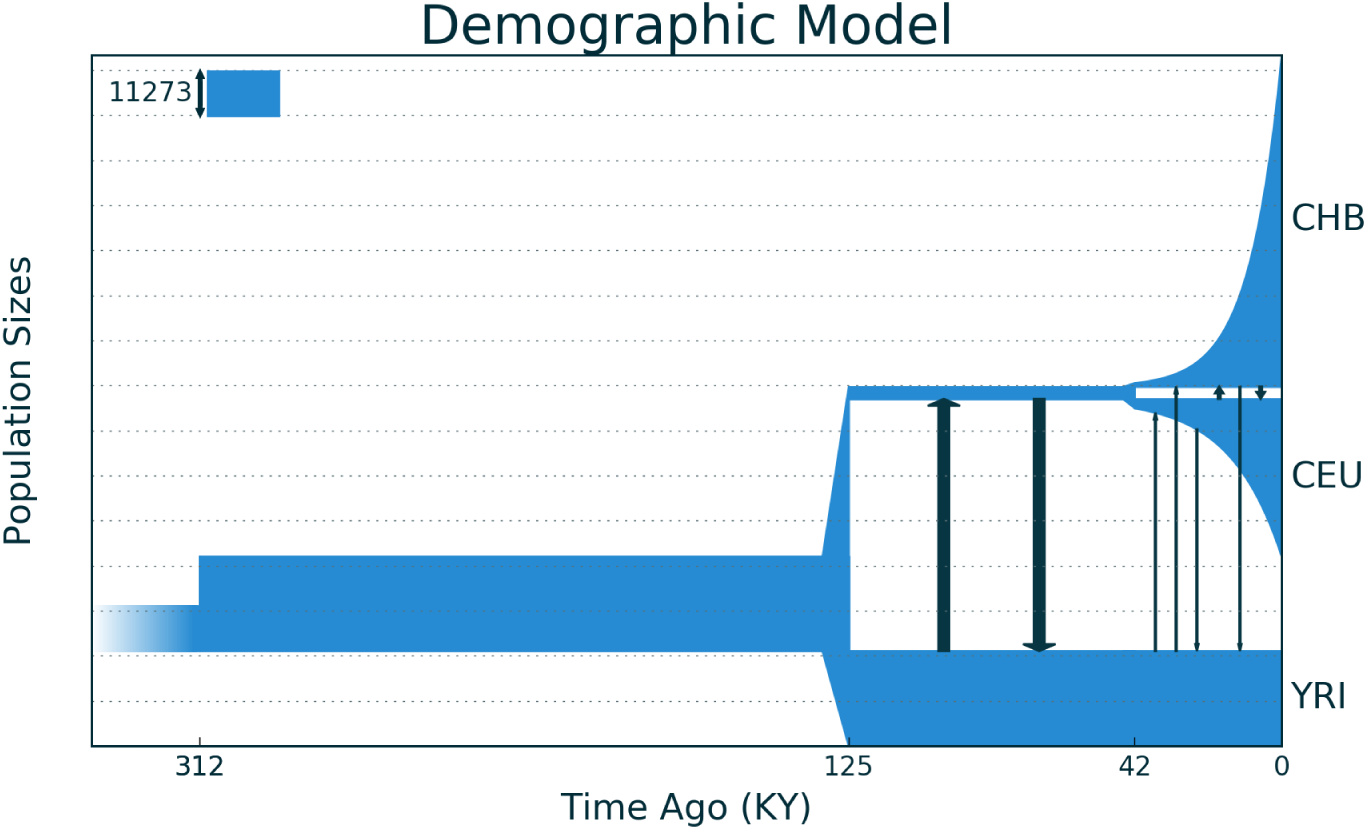
Out of Africa expansion.

We used a Powell optimizer to find maximum likelihood parameters (see Appendix F). Confidence intervals were obtained using recently introduced approaches for uncertainty estimation for composite likelihood [2]. Maximum-likelihood parameters are presented in Table 3.1.

**Table 2:** Parameters estimates inferred with the likelihood approach. For the mutation rate and the generation time, we used respectively: *μ* = 1.44 × 10^−8^ and *T_g_* = 29*y* [28]. We used 80 samples per population, performed the inference in genetic units, and scaled the parameters so that *N_A_* matches the point estimate from [9]. Con dence intervals are obtained by bootstrap over genic regions. Parameters are defined in Reference (??)

| Parameter | Estimate | 95% CI |
| --- | --- | --- |
| $N_A$ | 11273 | - |
| $N_{AF}$ | 23721 | 23033 - 24409 |
| $N_B$ | 3104 | 2882 - 3326 |
| $N_{EU0}$ | 2271 | 2091 - 2452 |
| $r_{EU}(\%)$ | 0.196 | 0.189 - 0.203 |
| $N_{AS0}$ | 924 | 864 - 984 |
| $r_{AS}(\%)$ | 0.309 | 0.30 - 0.318 |
| $m_{AF-B}(\times 10^{-5})$ | 15.8 | 14.7 - 16.8 |
| $m_{AF-EU}(\times 10^{-5})$ | 1.10 | 0.99 - 1.22 |
| $m_{AF-AS}(\times 10^{-5})$ | 0.48 | 0.42 - 0.55 |
| $m_{EU-AS}(\times 10^{-5})$ | 4.19 | 3.86 - 4.51 |
| $T_{AF}(kya)$ | 312 | 259 - 365 |
| $T_B(kya)$ | 125 | 110 - 139 |
| $T_{EU-AS}(kya)$ | 42.3 | 40.6 - 44.0 |

**Table 3:**
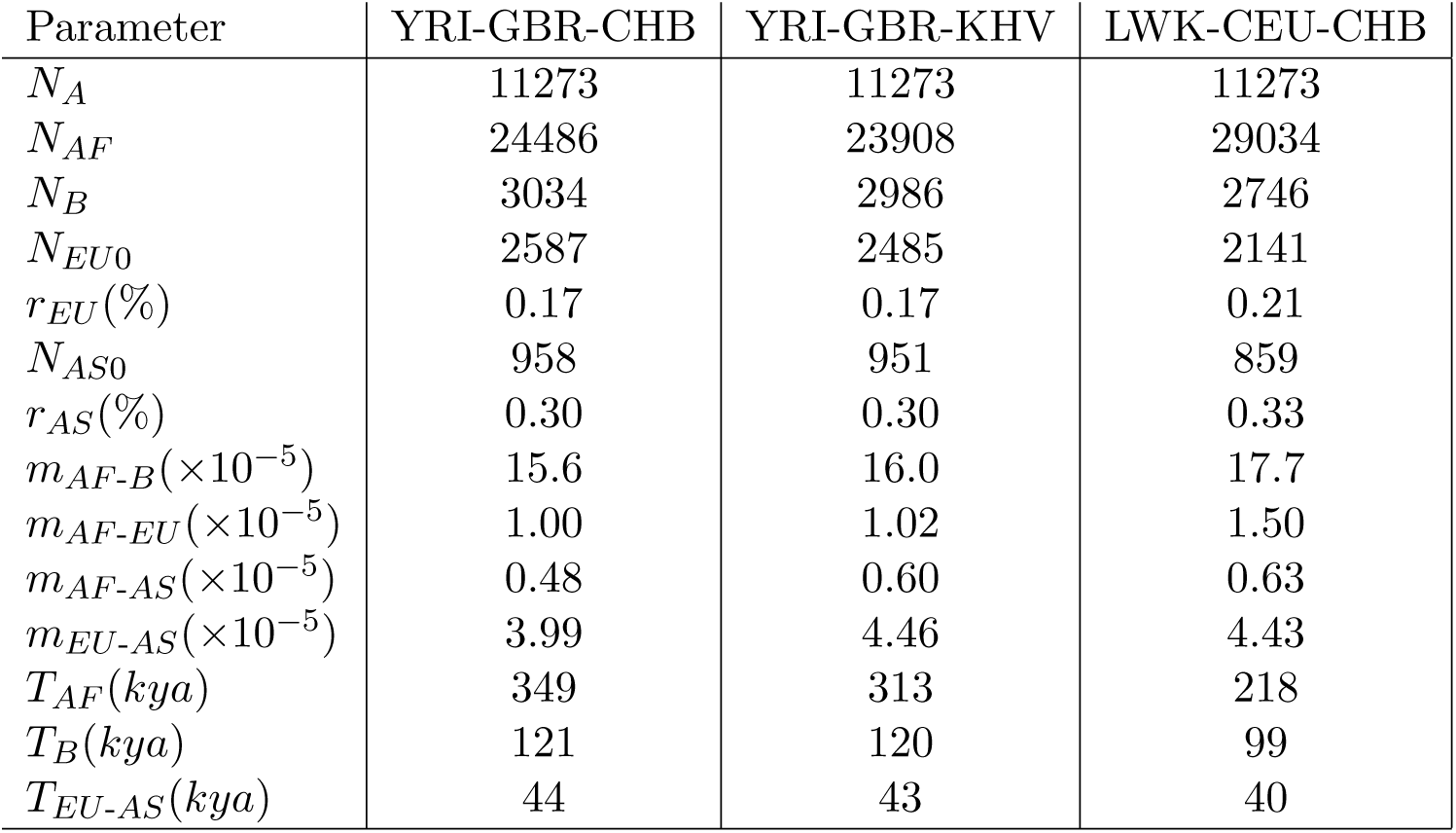
Parameters estimates inferred for the Out of Africa with different trios of african, european and asian populations from the 1000 Genomes dataset. For the mutation rate and the generation time, we used respectively: *μ* = 1.44 × 10^−8^ [11] and *T_g_* = 29*y* [28]. We used 80 samples per population. We performed the inference in genetic units, and scaled the parameters so that *N_A_* matches the point estimate from [9].

| Parameter | YRI-GBR-CHB | YRI-GBR-KHV | LWK-CEU-CHB |
| --- | --- | --- | --- |
| $N_A$ | 11273 | 11273 | 11273 |
| $N_{AF}$ | 24486 | 23908 | 29034 |
| $N_B$ | 3034 | 2986 | 2746 |
| $N_{EU0}$ | 2587 | 2485 | 2141 |
| $r_{EU}(\%)$ | 0.17 | 0.17 | 0.21 |
| $N_{AS0}$ | 958 | 951 | 859 |
| $r_{AS}(\%)$ | 0.30 | 0.30 | 0.33 |
| $m_{AF-B}(\times 10^{-5})$ | 15.6 | 16.0 | 17.7 |
| $m_{AF-EU}(\times 10^{-5})$ | 1.00 | 1.02 | 1.50 |
| $m_{AF-AS}(\times 10^{-5})$ | 0.48 | 0.60 | 0.63 |
| $m_{EU-AS}(\times 10^{-5})$ | 3.99 | 4.46 | 4.43 |
| $T_{AF}(kya)$ | 349 | 313 | 218 |
| $T_B(kya)$ | 121 | 120 | 99 |
| $T_{EU-AS}(kya)$ | 44 | 43 | 40 |

### 3.2 Validation with different trios of populations

Because of data and computational limitations, most previous studies have only considered a single triple of populations for estimating an Out-Of-Africa model. We wondered whether other populations would yield consistent results. Here we inferred the parameters with three different trios incorporating data from Luhya from Kenia (LWK), British (GBR) and Kinh Vietnamese (KHV) populations in addition to the three original populations we used. Inference results are presented in Table 3.2.

The inference is robust to permutation of European and Asian populations, in the sense that inferred parameters across triplets are consistent with the confidence intervals obtained in the YRI-CEU-CHB triplet. However, changing the African population from YRI to LWK changes parameters beyond the confidence intervals. This is because the reported confidence intervals take into account the finite nature of the genome, but not the variation across populations. Parameters inferred from different triplets are not wildly different, reflecting the shared or similar histories of some of the populations, but quantitative details can vary substantially. Perhaps the most interesting difference is the more recent inferred split between between LWK and Eurasians, compared to the split between YRI and Eurasians, supporting the idea that the YRI-Eurasian split time at 120 thousand years ago represents structure that persisted in Africa over tens of thousands of years before the Out-of-Africa event. This naturally begs for more detailed modelling of the relationships between African populations in the 1000 Genome project, but this requires a more thorough discussion of the historical and archeological context, and is outside the scope of this manuscript.

### 3.3 Out of Africa model with 4 and 5 populations

As *moments* handles up to five populations, we can go further and simulate more complex models than the Out of Africa model described in [12]. In this Section, we consider a model of Out of Africa expansion with four populations as we added a second asian population, the Japanese from Tokyo (JPT). The model is very similar to the original one described in Section 3.1 but we add a split in the Asian population resulting in the Chinese and Japanese populations (see Figure 2 and Table 3.3). We fixed the parameters inferred above in the three populations model and inferred the new parameters induced by the Asian split.

**Figure 2:**
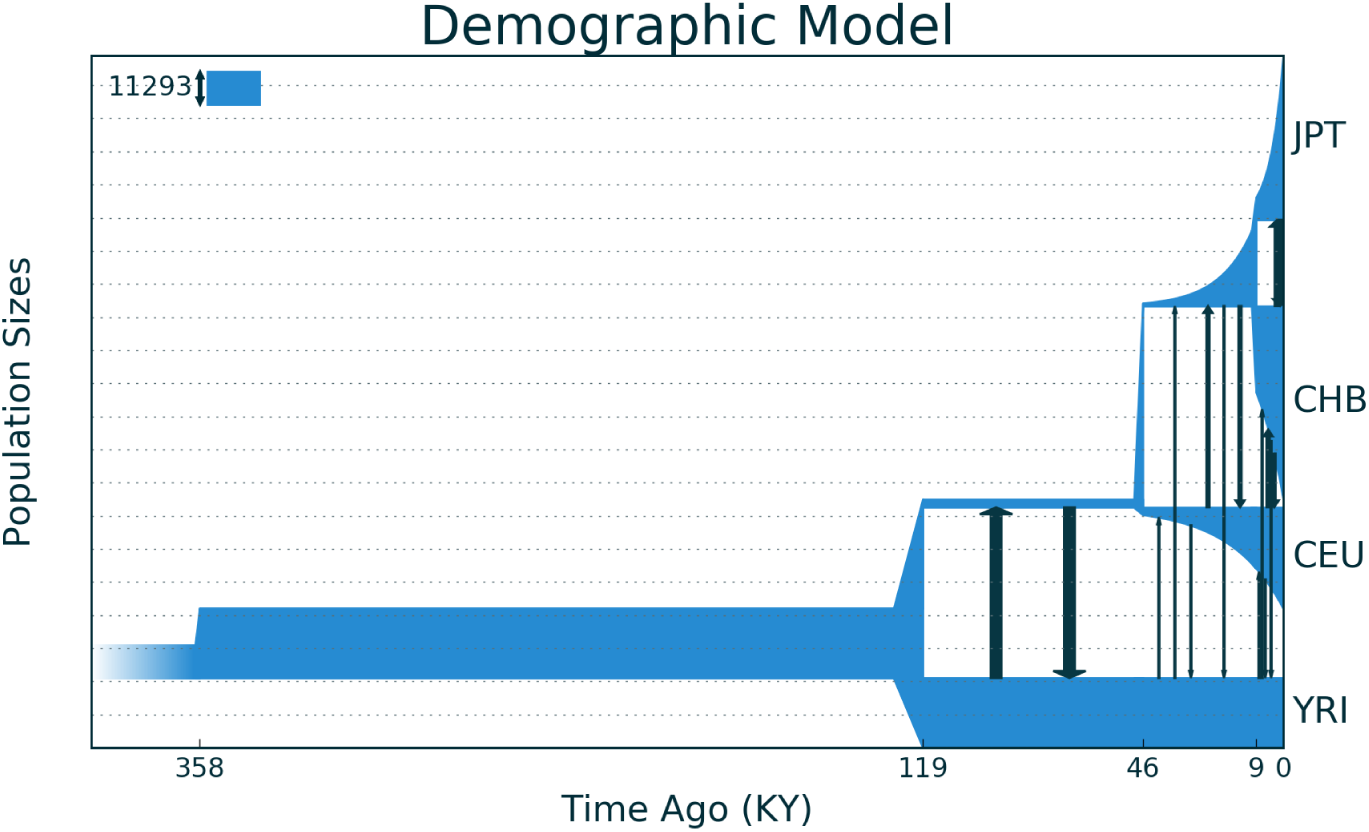
Out of Africa expansion model with four populations.

We also considered a model with five populations by adding a third Asian population, the Kinh Vietnamese (KHV). We fixed the parameters inferred from the four populations model and inferred the new parameters (see table 3.3) introduced by the split which gives rise to the KHV population.

## Discussion

We described a new approach to simulate the evolution of allele frequency distributions over time. On a practical level, this approach can be used wherever the diffusion approximation was applicable, in which case it typically provides faster, more accurate, and more robust solutions. Our software implementation will provide an easy transition to *∂a∂i* users: data handling and ancillary methods for *moments*are largely copied from open-source code from *∂a∂i*, model specification is simplified, and the number of adjustable parameters is reduced.

On a theoretical level, we proposed an exact and numerically robust solution to the discrete Wright-Fisher model. Even though its derivation is simple, we are not aware that such an approach has been used before. The exact approach is not practical under selection (where the moment equations does not close) and under migration (where the equations close only in a high-dimensional space). We bypass both difficulties by using a jackknife extrapolation inspired by capture-recapture models that is increasingly accurate for large sample size. This provides a simple, closed equation for the evolution of the allele frequency distribution that bypasses the diffusion approximation.

The applications of our solver to data from the 1000 Genomes project largely recapitulate and refine worldwide models of genetic diversity. Within Asia, we found that the best model involved a split between Han Chinese, Japanese, and Kinh Vietnamese approximately 9000 years ago. Importantly, we found the choice of a ‘representative’ population in Out-of-Africa models can substantially affect the inferred parameters, even ones not directly involving the population. Unsurprisingly, the choice of the African population has the largest impact on the inferred demography, emphasizing that previous OOA models may be applicable to many Eurasian populations, but not to other African populations. Building models including more than one African population will likely provide much more information about Human ancestry both in Africa and across the world.

Coalescent models are now extremely efficient at modelling neutral evolution, including with very large number of populations [14, 15]. One of the greatest advantages of forward approaches is the ability to handle a broader range of evolutionary scenarios. The moment approach proposed here broadens the range of scenarios where the allele frequency distribution can be modelled directly, including reversible mutations and selection. We expect that this will facilitate in particular the modelling of evolution over very long time-scales.

**Table 4:** Parameters estimates inferred with the likelihood approach for the 4 populations model. We used 30 samples per population.

| Parameter | Estimate |
| --- | --- |
| $N_A$ | 11293 |
| $N_{AF}$ | 23721 |
| $N_B$ | 2831 |
| $N_{EU0}$ | 2512 |
| $r_{EU}(\%)$ | 0.16 |
| $N_{AS0}$ | 1019 |
| $r_{AS}(\%)$ | 0.26 |
| $N_{JP0}$ | 4384 |
| $r_{JP}(\%)$ | 1.29 |
| $m_{AF-B}(\times 10^{-5})$ | 16.8 |
| $m_{AF-EU}(\times 10^{-5})$ | 1.14 |
| $m_{AF-AS}(\times 10^{-5})$ | 0.56 |
| $m_{EU-AS}(\times 10^{-5})$ | 4.75 |
| $m_{CH-JP}(\times 10^{-5})$ | 3.3 |
| $T_{AF}(kya)$ | 357 |
| $T_B(kya)$ | 119 |
| $T_{EU-AS}(kya)$ | 46 |
| $T_{CH-JP}(kya)$ | 9 |

**Table 5:** Parameters estimates inferred with the likelihood approach for the 5 populations model. We used 30 samples per population.

| Parameter | Estimate |
| --- | --- |
| $N_A$ | 11293 |
| $N_{AF}$ | 23721 |
| $N_B$ | 2831 |
| $N_{EU0}$ | 2512 |
| $r_{EU}(\%)$ | 0.16 |
| $N_{AS0}$ | 1019 |
| $r_{AS}(\%)$ | 0.26 |
| $N_{KHV0}$ | 2356 |
| $r_{KHV}(\%)$ | 17.4 |
| $N_{JP0}$ | 4384 |
| $r_{JP}(\%)$ | 1.29 |
| $m_{AF-B}(\times 10^{-5})$ | 16.8 |
| $m_{AF-EU}(\times 10^{-5})$ | 1.14 |
| $m_{AF-AS}(\times 10^{-5})$ | 0.56 |
| $m_{EU-AS}(\times 10^{-5})$ | 4.75 |
| $m_{CH-KHV}(\times 10^{-5})$ | 21.3 |
| $m_{CH-JP}(\times 10^{-5})$ | 3.3 |
| $T_{AF}(kya)$ | 357 |
| $T_B(kya)$ | 119 |
| $T_{EU-AS}(kya)$ | 46 |
| $T_{CH-KHV}(kya)$ | 9.8 |
| $T_{CH-JP}(kya)$ | 9 |

## A Coefficients of the discrete operators

In this section we first present the full form of the discrete drift, selection, and migration operators involved in equation (7) in the simplest model where selection, migration, and drift are treated linearly, and then present an outline of the derivation.

Remember that the ODE for an AFS in *p* populations, with *N_j_* diploid individuals in population *j* and *n_j_* sequenced ploids per population, is

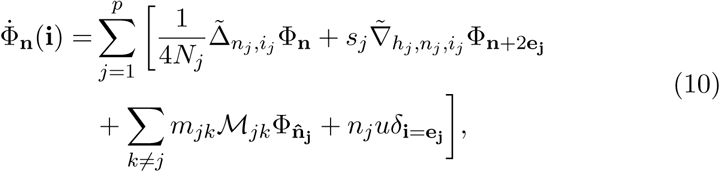
where 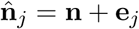 is the parental sample size required to reach sample size **n** after migration from *j* to *k*, and *e_j_* is the *j^th^* vector of the canonical basis of 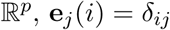.

The drift operator 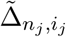 has the form

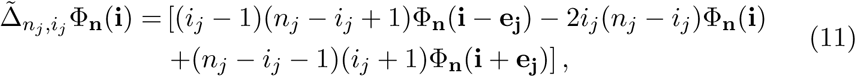

and is closely related to transition matrices derived in [5, 26]. The selection operator 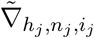 is defined by

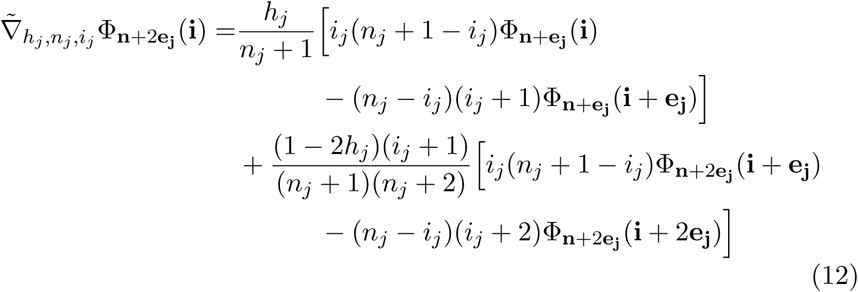

and the migration operator *ℳ_jk_* is

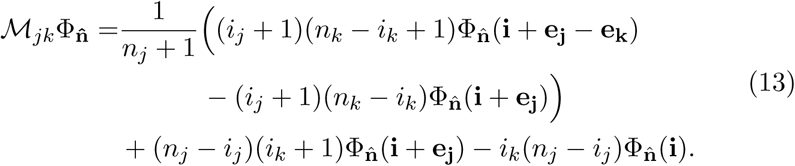

### A.1 Derivation for the drift operator

For simplicity, we derive the drift and selection operators in one dimension, since the multi-dimension case is a straightforward generalization with a heavier notation burden. The probability of observing a single two-way coalescence in a sample of size *n* is

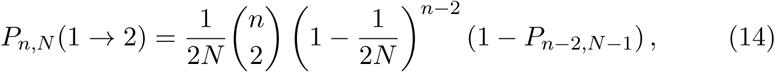
where 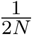 is the probability that a given pair of lineages coalesces, 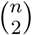 is the number of distinct pairs of lineages that can coalesce, 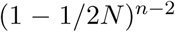 is the probability that there is no additional coalescence to the same ploid, and 1 − *P*_*n*−2,*N*−1_ is the probability that none of the remaining *n* − 2 lineage coalesce to the remaining *N* − 1 ancestors. If we only keep leading order terms in 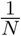, we find the classical large-population limit

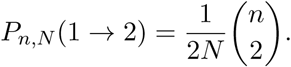

Given a single two-way coalescence, we then want to compute the frequency distribution in the offspring given the distribution in the parental generation. We imagine that the *n* descendent ploids are copied from *n* parental ploids, with one parental ploid drawn twice, and one parental ploid never drawn. Since the *n* parental ploids were drawn independently, their expected allele frequency distribution corresponds to the expected frequency in the parental generation. To compute the change in expected allele frequency between parental and offspring generation, we must compute the probability of allele frequency changes between parents and offsprings.

The probability of observing a transition from parental frequency *i* + 1 to descendent frequency *i* after a two-way coalescence is the probability that the twice-selected ploid carries the reference genotype (which we represent as 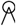), and the unselected ploid is non-reference (which we represent as •):

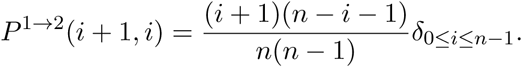

Similarly, the gain of an alternate ploid results from event 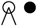 and has probability

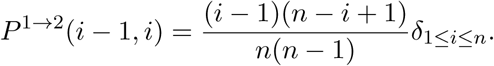

The probability of transitioning from *i* alternate ploids to *i* − 1 (via 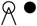) or *i* + 1 (via *i* + 1 (via 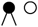) is

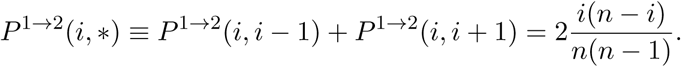

We can finally compute the drift operator for a single two-way coalescence:

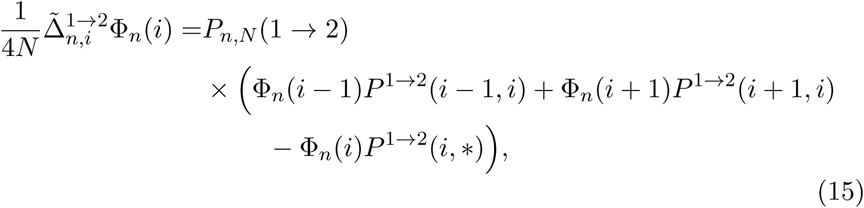
which simplies to Equation (11).

### A.2 Derivation of the selection operator

Here we derive a one-dimensional version of Equation (12). The derivation is elementary but a bit cumbersome. See Appendix C for a different derivation that relies on the diffusion approximation.

We consider the action of selection on a single transmission, when a new ploid is drawn from a diploid individual. We derive the results in the context of negative selection (*s* < 0 and *sh* < 0) where we can think of selection as eliminating a proportion of the neutral transmissions. A transmission is eliminated by selection with probability −*sh* if the parent is a heterozygote, and with probability −*s* if the parent is an alternate homozygote. For bookkeeping, we write the state of the parent as 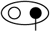, with the empty circle representing a reference allele, solid circle representing the alternate allele, and the vertical line representing the putative initial selected ploid. When selection acts, this ploid is replaced by a new ploid, taken at random from the parental population. Since we assume that at most one selective event occurs per generation, we do not need to know the diploid state of this replacement ploid. In the selective event labeled as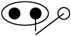, the transmission of an alternate allele from a homozygous ancestor was eliminated and replaced by transmission from a reference allele. To generate n descendent ploids with one selective event, there have been a total of *n* + 2 relevant ploids: the *n* selected initially, the diploid companion of the rejected ploid and the replacement ploid. We want to compute the change in allele frequency caused by the selection process relative to the neutral transmission of the n selected ploids. We write 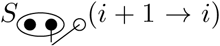 for the probability that the *n* initial parental ploids included *i* + 1 alternate alleles, but the offspring ploids include *i* because of selective process 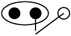. This is proportional to the number Φ_*n*+2_(*i* + 2) of loci having the required *i* + 2 alternate alleles in a sample of size *n* + 2, the probability of selecting the correct triplet of ploids (namely the hypergeometric 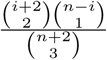 for the correct choice of three ploids times 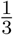 for the correct order), the probability that the selection event occurs for one transmission (−*s*), and the number of transmissions where selection can act (*n*). Putting all these together, we get

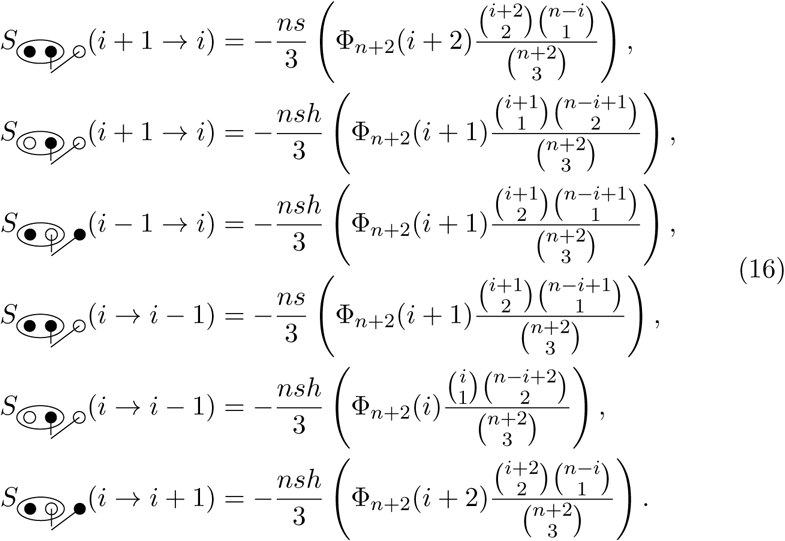

The first three terms, of the form *S* (· → *i*), describe increases to the number of alleles at frequency *i*. The last three terms, of the form *S* (*i* → ·); contribute to a decrease relative to the neutral case: Their contribution to the evolution equation will have opposite signs.

Under an additive model, the diploid companion to the selected allele plays no role and the evolution should only depend on Φ_*n*+1_ rather than Φ_*n*+2_. To make this explicit, we collect terms into an additive term, proportional to *h*, and a dominance term, proportional to (1 − 2*h*). The coefficient of the dominance term can be obtained by setting *h* = 0 in the list of contributions from Equation (16). The dominance term reads

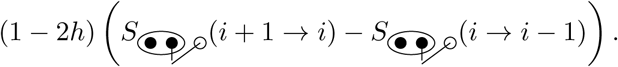

The additive term contains the remaining terms

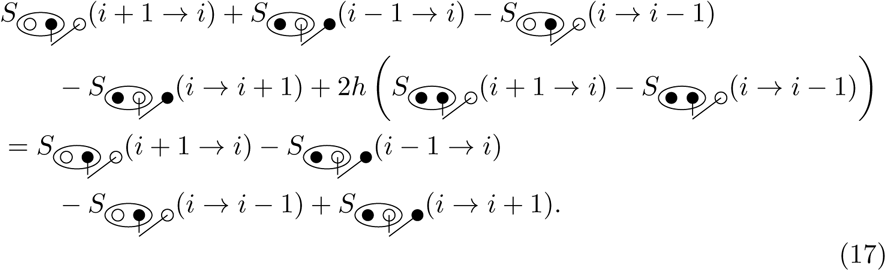

We combine the first and last term using the downsampling formula

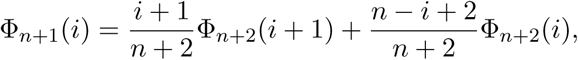

and similarly combine the middle two terms using

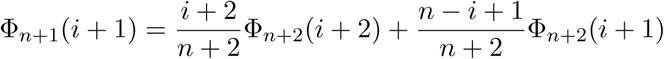

to reach Equation (12).

### A.3 Derivation of the migration operator

We derive the migration operator for populations *j* and *k*, with *m_jk_* the probability that a ploid in population *k* has a parent from population *j*. We consider a sample of *n_j_* ploids from population 1 and *n_k_* from population *k* and suppose that the migration rate is small enough that at most one migration occurs per site per generation.

In this case there are only two ploids involved in the migration, the replaced and the replacement ploids, and two possible configurations that result in changes in the allele frequency, 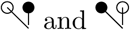, where circles represent ploids: a dark circle represents an alternate allele, and the ploid on the left is the migrant replacing the ploid to the right. If 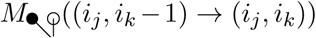 is the rate of increase of the number of loci with alternate allele counts (*i_j_*, *i_k_*) because of process 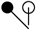 we have two rates of increase:

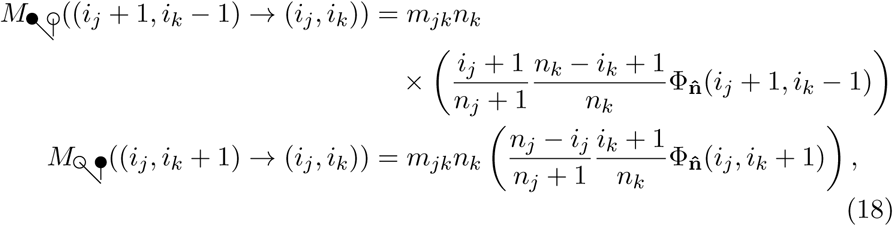

and two corresponding rates of decrease:

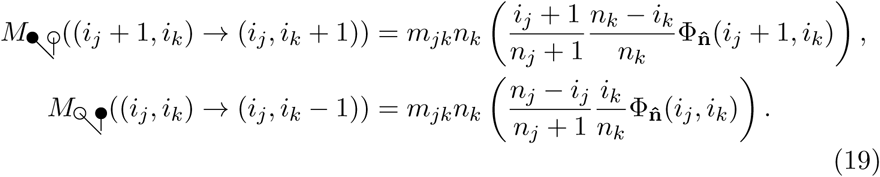

We can combine these to get the transition rate due to migration:

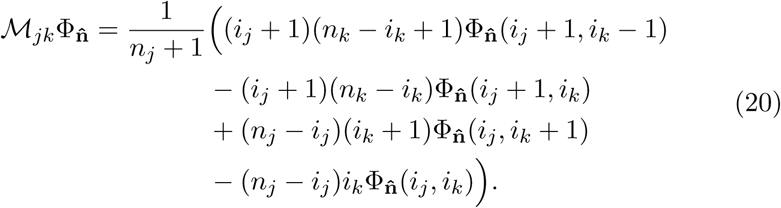

The code in *moments* uses a slightly different equation that can be derived from this by the application of the downsampling formulas, namely

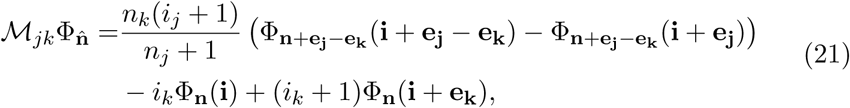

This makes it possible to only use the jackknife on the first two terms.

## B Generalized forms of the operators

### B.1 Fifinite genome model for mutations

To facilitate comparison with previous work, we focused in this work on the infinite-sites model, assuming that mutations occur at previously invariant loci and neglecting back mutations. However, finite genome and back mutations are easily accommodated by the present model. In this case, the single population mutation term *M_i_*Φ_*n*_ is:

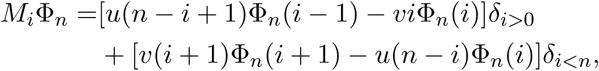

where *u* and *v* are the forward and backward mutation rates.

### B.2 Drift with multiple coalescences

The standard diffusion approximation and Kingman’s coalescent model neglect the possibility that multiple coalescences may occur during the same generation in the history of a sample. We used a similar approximation to compute the drift operator in Appendix A. Even though multiple coalescences are indeed rare for small sample sizes and large populations, they can have a measurable effect for large sample sizes [14].

Here we consider the next-order correction to the drift operator by accounting for three-way and double two-way coalescences (Figure 3), each contributing corrections of order 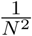.

**Figure 3:**
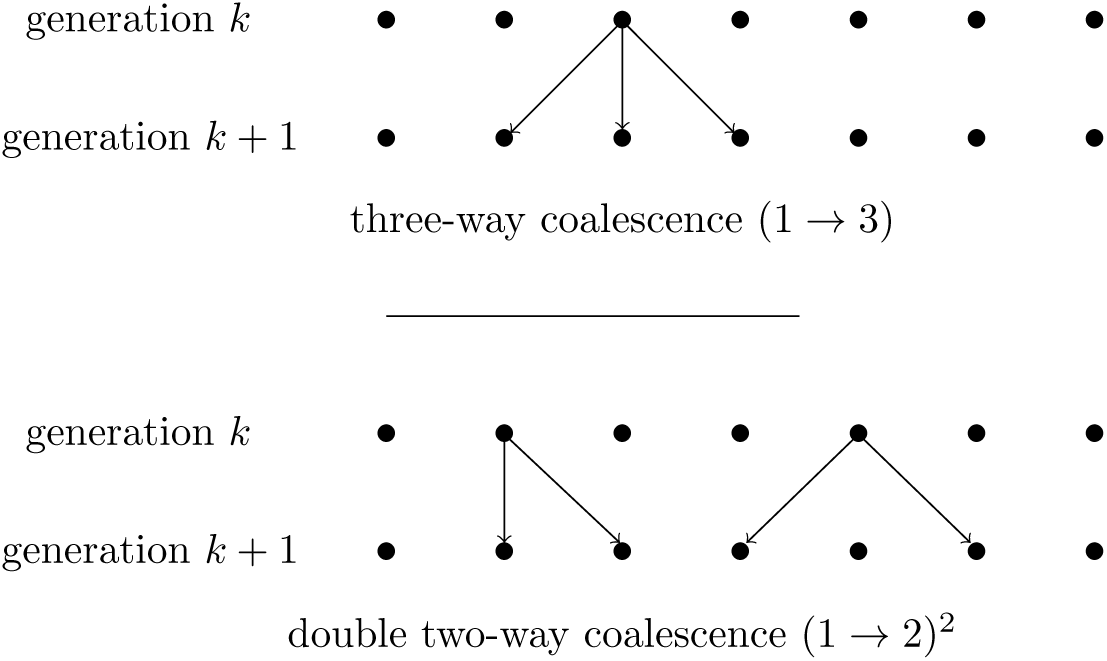
Multiple coalescences.

We can decompose the drift operator ∆_*n,i*_ in contributions from two-way, three-way, and double two-way coalescence

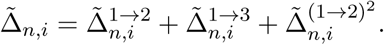

The computation of individual terms, outlined below, is cumbersome but elementary. The key result is that the rate of change in Φ_*n*_(*i*) is a linear combination of the Φ_*n*_(*j*) with *j* ranging from *i* − 2 to *i* + 2.

#### B.2.1 Single two-way coalescent

The two-way coalescent contribution is also described by Equation (15), but if we consider multiple coalescences we must also account for corrections to *P*_*n,N*_ (1 → 2) of order 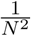, i.e.,

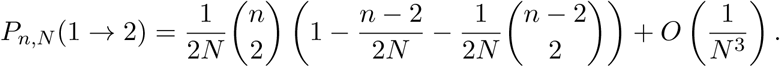

#### B.2.2 Single three-way coalescent

Similarly, the probability of a three-way coalescent is, to leading order in *N*, is

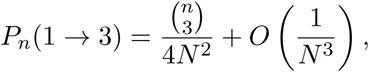

and we must now compute 5 transition probabilities.

For example the transition probability from *i* + 2 to *i* is,

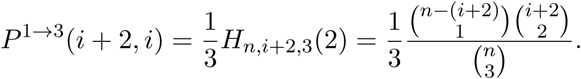

Here 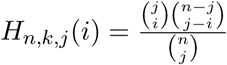 is the hypergeometric distribution with *j* trials and *i* successes, sampling from a finite population of size *n* with *k* successes. To derive this result, we note that a triple coalescence can create a transition from *i* + 2 to *i* if it involves three parental ploids: one (reference) chosen three times, and two (alternate) never chosen: 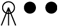. The probability of this happening is the product of the probability of drawing two alternate ploids in a sample of three (i.e., *H*_*n,i*+2,3_(2)), times the probability that we picked the reference allele to coalesce (i.e., 1/3).

Similarly, we have contributions from 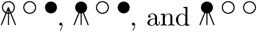

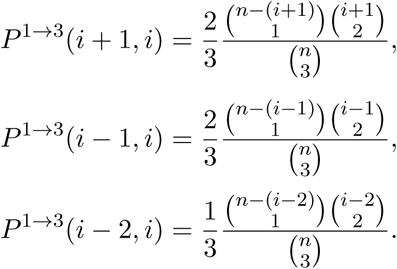

Finally we want the probability of changing from ancestral frequency *i* to any other offspring frequency. Because the only way to maintain the number of reference ploids under a triple coalescence is to pick either three reference ploids 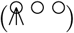 or three non-reference ploids 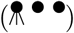, we have

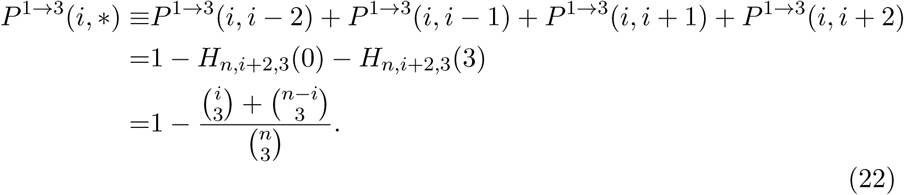

We finally have the corresponding drift operator

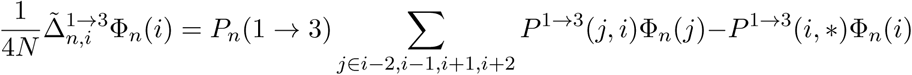

#### B.2.3 Double two-way coalescent

The probability of drawing a double coalescence is

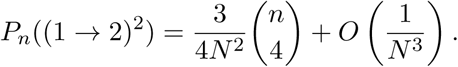

Here we need to consider four parental ploids with two coalescences and two not selected. We have a term corresponding to 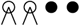

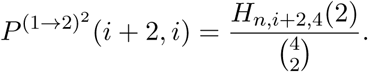

Similarly 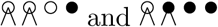 contribute to

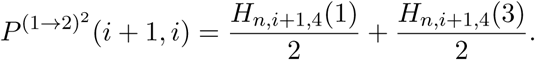

We then have the complementary contributions 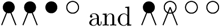 to

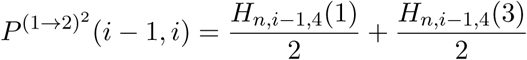

and 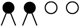 for

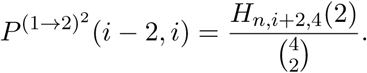

There are now three possibilities for not changing the allele frequency, namely 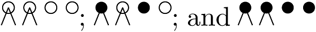, so that the probability of changing from frequency *i* given a double two-way coalescence is

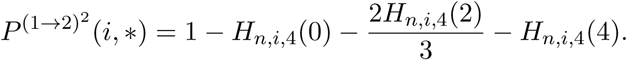

The drift operator is therefore

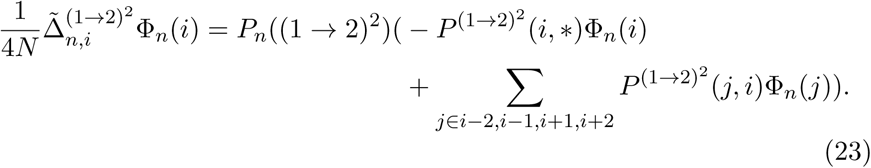

Changes to Φ_*n*_(*i*) caused by drift are a simple linear function of the Φ_*n*_(*j*) fir *j* ∈ *i* − 2,…, *i* + 2. This operator is sparse for large n and easy to compute numerically. Higher-order corrections in 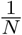 would simply involve more terms and a progressively denser linear operator.

### B.3 Strong migration and admixture

To compute the frequency spectrum 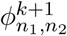 after admixture, we need to consider the origin of *n*_1_ + *n*_2_ lineages. Under strong recent bidirectional migration, there are loci at which all *n*_1_ + *n*_2_ lineages come from population 1, and other loci at which all lineages come from population 2. Thus 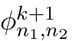 depends on the set 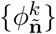 for all **ñ** containing *n*_1_ + *n*_2_ lineages. This information is contained in the frequency spectrum 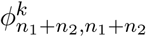.

Specifically, we can write

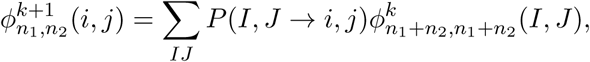

Where *P*(*I*, *J* → *i*, *j*) is the probability of observing alternate allele counts (*i*, *j*) when each allele is drawn independently and without replacement from the two ancestral samples with frequency (*I*, *J*). The direct computation of *P*(*I*, *J* → *i*, *j*), summing over the possible inheritance patterns, is straightforward but computationally demanding for large sample sizes or high-dimensional applications.

Instead, we used a dynamic programming approach that is a bit technical but allows for important speedups. The general idea is that we perform migrations one ploid at a time, rather than simultaneously, allowing successive migrants to replace previous ones. The end result is not quite the distribution that we want, because the bulk replacement process does not allow for replacement among migrants. However, we can easily compute and correct for this difference. The main benefit of this approach is that we avoid having to perform multiple integrals over the possible migrant configurations.

To simulate one-way admixture from population 1 into population 2 at rate *m*_12_, we therefore first allow one lineage at a time from population 1 to migrate into population 2. We sequentially compute the spectra *ψ_R_* resulting from *R* = 0, 1 2,…, *v* migrating lineages, with chosen so that all plausible migrant numbers are covered (details below). The *ψ_R_* do not correspond to the correct frequency distribution, because the number of migrant alleles per site does not have the correct distribution: Our goal is then to choose a linear combination 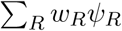 such that

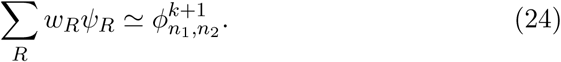

In the standard infinite-sites model, the number of replaced lineages in the admixed sample follows the binomial distribution *B*(*n*, *m*_12_). We can therefore write the frequency spectrum 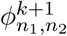 as the sum 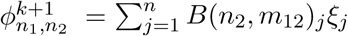, where *ξ_j_* is the frequency spectrum that would result if exactly *j* lineages were replaced through migration.

To identify the set of *w_R_* that satisfy Equation (24), we also express *ψ_R_* as a linear combination of the *ξ_j_*. The probability Γ_*jR*_ of observing *j* net replacements after *R* sequential replacements is easily computed numerically. This leaves us with 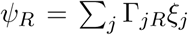. If we express both sides of Equation (24) in terms of the *ξ_j_*, we find

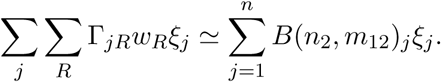

We would then like to choose the *w_R_* such that the coeffcients are equal:

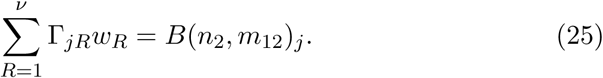

Provided that *v* is large enough, this linear problem admits solutions for *w_R_*. Unfortunately, exact solutions to this equation are prone to strong oscillations in the *w_R_*, which lead to numerical instabilities. Rather than seeking exact but oscillating solutions to Equation (25), we seek non-negative solutions that minimize the squared error 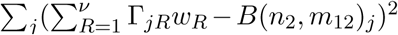 using the active set method implemented in the scipy.optimize.nnls routine. Assuming that we found a solution with an acceptably small error, we finally use Equation (24) to compute the frequency spectrum.

To choose *v*, we note that *v* sequential replacements give an average of approximately *n*(1 − *e^v/n^*) net replacements once multiple replacements of the same lineage have been accounted for (e.g., [?]). We find that we get accurate results if we have enough net replaced lineages to cover most likely cases in binomial sampling: 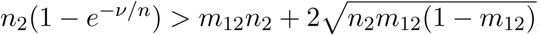.

## C ODEs on the moments

### C.1 One dimensional case

In this section, we show how the system of ordinary differential equations (7) derived in the main text can also be derived by integration by parts from the diffusion approximation to the Wright-Fisher process. Under the di u-sion approximation, the evolution of the single-population allele frequencies density *ϕ*(*x*, *t*) follows (e.g., [12]):

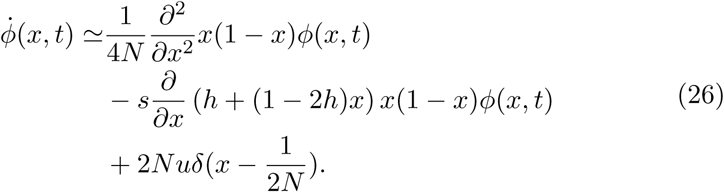

The first term of the right hand side is a diffusion term modeling the effect of genetic drift and the second term is a transport term that accounts for selection. Finally, the source term 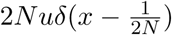 models the mutation process under the infifinite site assumption.

We are interested in the moment-like statistics Φ_*n*_(*i*), obtained projecting the allele frequencies density *ϕ* on the weight functions defined by 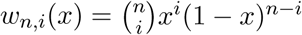:

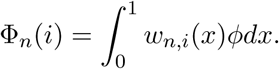

The time derivative of Φ_*n*_(*i*) is given by:

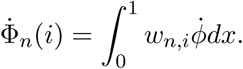

The delta function source term in Equation (26) is integrated easily. The leading-order term in 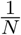 is simply *nuδ*_*i*=1_: this is the rate at which mutations appear on *n* lineages over one generations:

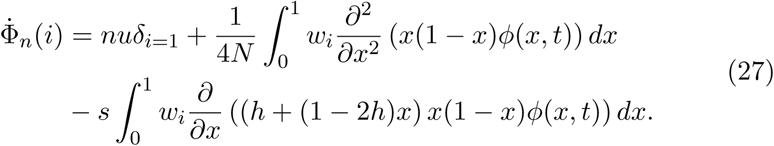

**drift term:** We want to rewrite the drift term of equation (27) in terms of the moments Φ_*n*_(*i*). To this end, we integrate by parts:

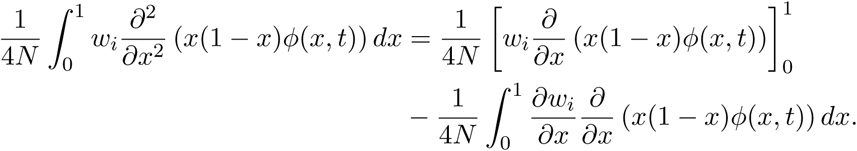

As *w_i_*(0) = *w_i_*(1) = 0 we assume that the first term is 0. We can integrate by parts the second term:

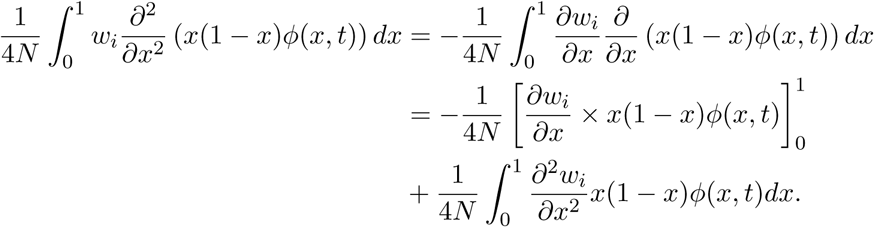

As we consider a finite genome, *ϕ*(0, *t*) and *ϕ*(1, *t*) are finite and, once again, the term in square brackets is zero. Moreover, we have:

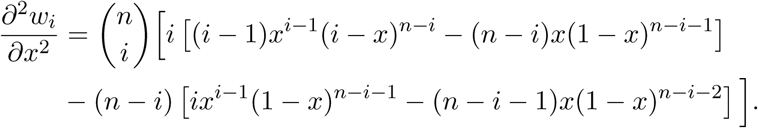

Thus,

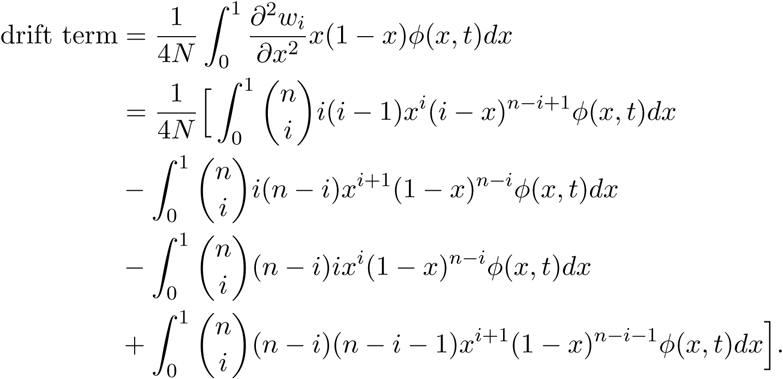

Rearranging this expression, we can write it in terms of the Φ_*n*_(*i*):

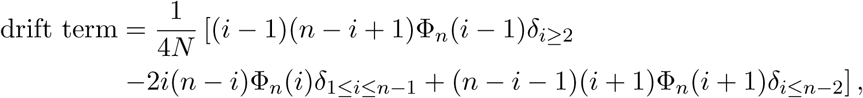
which corresponds to the one-dimensional version of equation (11).

**selection term:** Here again, we integrate by parts the selection term:

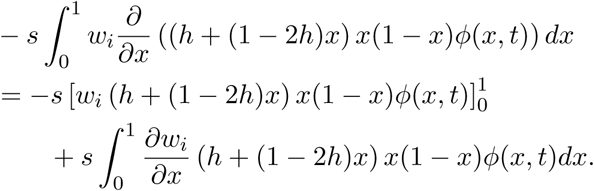

We can again assume that the term in square brackets is zero, we rearrange the other integral term in order to write it in terms of the Φ_*n*_(*i*). We finally get:

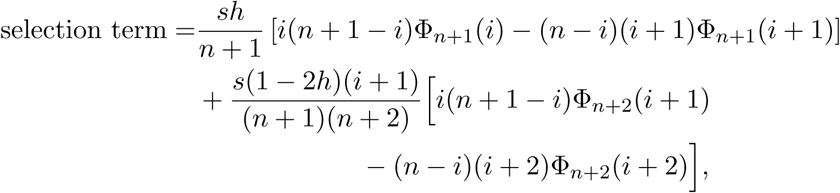

which corresponds to the one-dimensional version of equation (12).

Bringing these expressions together, we get a system of ordinary differential equations on the Φ_*n*_(*i*):

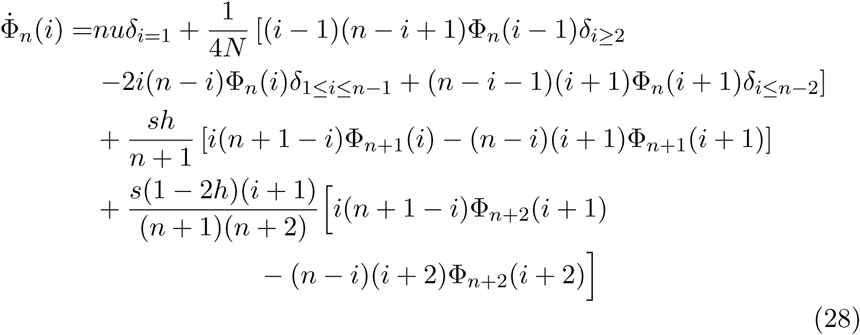

This is the one-dimensional equivalent to equation (10).

### C.2 Extension to multiple dimensions

The diffusion model can be generalized to multiple population studies. The drift, selection and mutation terms are directly derived from the unidimensional case. We just need to add the effect of the migrations between the populations. The equation on the joint frequency spectrum is:

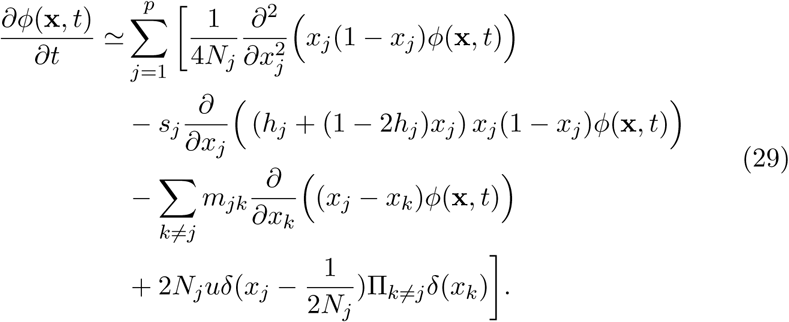

Now **x** is a vector: **x** = [*x*_1_,…, *x*_p_] ∈ [0; 1]^p^. It is the same for the parameters **s** and **h** and *N_j_* is the reference population size for population *j*. The term 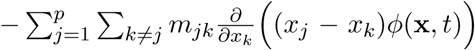 accounts for the migrations where *m_jk_* is the proportion of individuals in population *k* whose parents were born in population *j*.

We are interested in the multi dimensional statistics generalizing the Φ_*n*_(*i*):

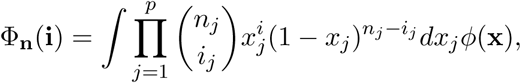

where *n* and *i* are vectors. The calculations are a bit more tedious but we can do the same as for the single population case: integrating by parts each term and writing it in terms of the Φ_*n*_(*i*), we recover Equation (10).

## D Jackknife approximation for moment closure

The evolution equations for Φ_*n*_ involves higher order terms, such as Φ_*n*+1_ or Φ_*n*+2_, which leads to a moment closure problem. To tackle this, we use a jackknife approach to approximate higher-order terms as linear functions of Φ_*n*_:

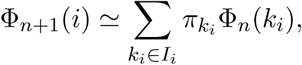

where *I_i_* is a set of indices chosen so that the frequency *k_i_/n* is close to the target frequency *i*/(*n* + 1):

Given a set *I_i_*, we choose the coeffcients *π_k_i__* so that the jackknife is exact for a given parameterized family of functions Φ_*n*+1_.

The ‘order’ of the jackknife refers to the number of terms in *I_i_*: the more terms, the more general the family of functions can be, but the more numerically unstable the jackknife becomes.

We focus here on the order 3 jackknife for the terms Φ_*n*+1_(*i*) and we will use the following approximation:

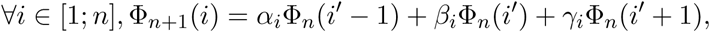

Where the index *i´* is chosen so that the frequency 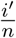 is as close as possible to 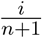 and to satisfy the following boundary conditions:

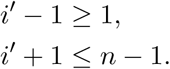

We choose the approximation so that the interpolation is exact for quadratic allele frequency distribution:

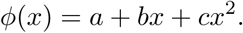

Even though this seems like a drastic approximation, the jackknife only requires the quadratic approximation to hold locally: the parameters *a*, *b*, and *c* will be chosen independently for each value of *i*.

The moments become:

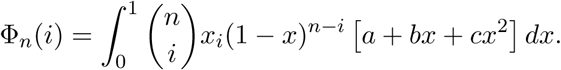

The integral can be performed analytically to yield:

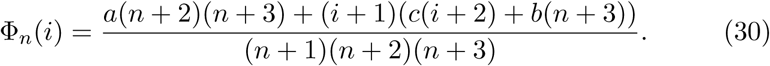

We then look for the jackknife coefficients *α_i_*, *β_i_* and *γ_i_* that cancel the approximation error:

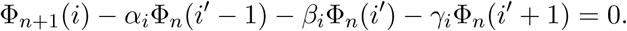

We replace the Φ by their expressions under Equation (30):

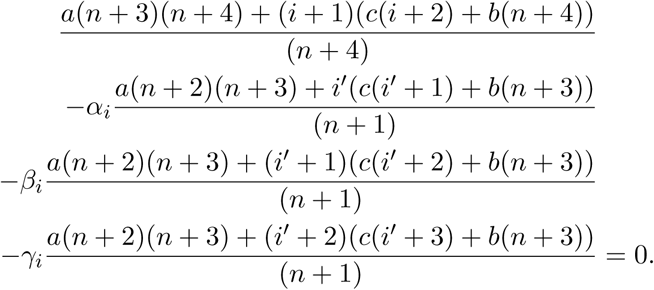

This equation should hold for all values of (*a*, *b*, *c*), so we can use the particular values (1, 0, 0), (0, 1, 0) and (0, 0, 1) and write the corresponding system of equations:

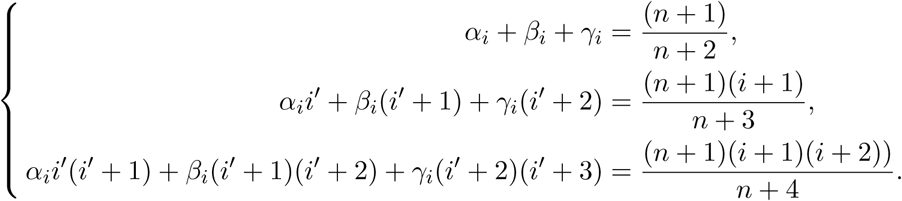

We can compute the jackknife coeffcients *α_i_*, *β_i_* and *γ_i_* by solving the previous linear system. We finally get:

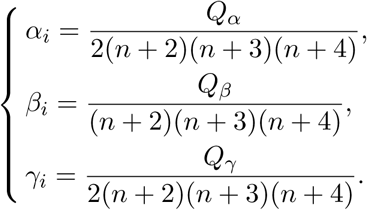

with

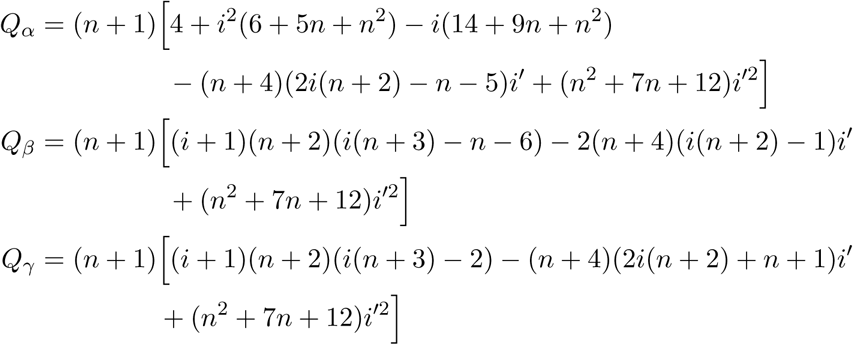

The same strategy is used to approximate Φ_*n*+2_(*i*) as a linear combination of the Φ_*n*_ entries. We use the same formulation for multidimensional simulations as the moment closure problem can be addressed separately in each dimension. For instance, in the 2-dimensional case, the selection in the first population will involve the higher order term Φ_*n*_1_+1,*n*_2__ (*i*_1_, *i*_2_) that we will approximate as follows:

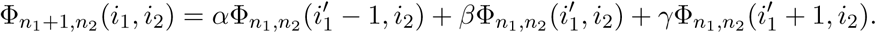

## E Benchmarks

In this section we present an extensive set of test cases to compare computational performances of *Moments* and *∂a∂i* in terms of speed and accuracy. Refer to paragraph 2.7 for more details about the metrics used for the comparisons. The results on test cases with 30, 80 and 200 ploids are respectively given in Tables 6, 7 and 8. A more explicit description of the test cases is given in Table 9. Unless otherwise specified, populations sizes grow linearly: *N*(*t*) = 1 + 0.01*t* with *t* the time in genetic units. For the exponential growth, we used the function *N*(*t*) = 10^*t*^. When dealing with several populations, we used the same selection and migration parameters as well as the same population growth functions for all the populations, except for the Out-of-Africa model.

**Table 6:** Performance comparisons between *∂a∂i* and Moments on several scenarios (30 samples per population). For *∂a∂i* simulations we used Richardson extrapolation to improve convergence. The time T provided is the simulation time in genetic units.

| Demographic model | method | exec time (s) | mean($\varepsilon_r$) | $\Delta$ LL | < 0 entries |
| --- | --- | --- | --- | --- | --- |
| Neutral equilibrium 1D | $\partial a \partial i$ | 3.64e-02 | 1.25e-02 | **1.47e-05** | 0 |
|  | *Moments* | **1.54e-03** | **9.74e-03** | 4.64e-05 | 0 |
| Neutral 1D,<br>T = 1.0 | $\partial a \partial i$ | 3.59e-02 | **3.71e-03** | **4.15e-06** | 0 |
|  | *Moments* | **1.20e-03** | 1.44e-02 | 3.24e-04 | 0 |
| Neutral 1D,<br>T = 5.0 | $\partial a \partial i$ | 1.42e-01 | 2.50e-03 | 1.17e-05 | 0 |
|  | *Moments* | **4.63e-03** | **2.82e-04** | **4.78e-08** | 0 |
| Selection 1D,<br>T = 1.0 | $\partial a \partial i$ | 3.70e-02 | **6.50e-03** | **1.13e-06** | 0 |
|  | *Moments* | **8.22e-03** | 1.45e-02 | 3.68e-04 | 0 |
| Selection 1D,<br>T = 5.0 | $\partial a \partial i$ | 1.48e-01 | 5.62e-03 | 3.42e-06 | 0 |
|  | *Moments* | **3.88e-02** | **9.23e-04** | **5.51e-07** | 0 |
| Neutral equilibrium 2D | $\partial a \partial i$ | 2.76e-01 | 1.25e-02 | 3.06e-03 | 0 |
|  | *Moments* | **4.16e-02** | **9.74e-03** | **9.29e-05** | 0 |
| Selection 2D,<br>T = 1.0 | $\partial a \partial i$ | 1.68e-01 | 1.25e-02 | 2.85e-03 | 0 |
|  | *Moments* | **2.12e-02** | **9.11e-04** | **-1.39e-03** | 0 |
| Selection 2D,<br>T = 5.0 | $\partial a \partial i$ | 6.20e-01 | 5.45e-03 | 9.26e-03 | 0 |
|  | *Moments* | **7.50e-02** | **9.18e-04** | **-6.54e-03** | 0 |
| Selection 2D,<br>T = 1.0, neutral fs0 | $\partial a \partial i$ | 1.68e-01 | **5.45e-03** | 4.72e-04 | 0 |
|  | *Moments* | **1.70e-02** | 1.73e-02 | **2.57e-04** | 0 |
| Selection, migration 2D,<br>T = 1.0 | $\partial a \partial i$ | 9.33e-01 | 8.13e-03 | 9.96e-05 | 0 |
|  | *Moments* | **2.20e-01** | **3.71e-03** | **2.11e-05** | 0 |
| Selection, migration 2D,<br>T = 5.0 | $\partial a \partial i$ | 4.44e+00 | 8.37e-03 | 4.07e-04 | 0 |
|  | *Moments* | **9.95e-01** | **2.53e-03** | **2.07e-05** | 0 |
| Selection, migration 2D,<br>T = 1.0, neutral fs0 | $\partial a \partial i$ | 9.14e-01 | 5.22e-03 | 1.33e-04 | 0 |
|  | *Moments* | **2.18e-01** | **1.49e-03** | **1.49e-05** | 0 |
| Fast growth 2D,<br>T = 1.0 | $\partial a \partial i$ | 9.61e-01 | 2.61e-01 | **7.06e-04** | 16 |
|  | *Moments* | **2.20e-01** | **6.29e-02** | 7.96e-04 | 0 |
| YRI-CEU 2D | $\partial a \partial i$ | **1.30e-01** | **3.64e-03** | **3.43e-05** | 0 |
|  | *Moments* | 1.89e-01 | 8.25e-03 | 1.49e-04 | 0 |
| Neutral equilibrium 3D | $\partial a \partial i$ | 1.58e+01 | 1.25e-02 | 9.14e-03 | 0 |
|  | *Moments* | **4.69e-01** | **9.74e-03** | **1.39e-04** | 0 |
| Selection 3D,<br>T = 1.0 | $\partial a \partial i$ | 6.58e+00 | 7.37e-01 | 1.77e-03 | 0 |
|  | *Moments* | **1.21e-01** | **4.44e-05** | **-5.32e-02** | 0 |
| Selection 3D,<br>T = 5.0 | $\partial a \partial i$ | 2.70e+01 | 7.38e-01 | 6.16e-03 | 0 |
|  | *Moments* | **5.75e-01** | **8.82e-06** | **-1.74e-01** | 0 |
| Selection 3D,<br>T = 1.0, neutral fs0 | $\partial a \partial i$ | 6.28e+00 | **3.62e-03** | 1.77e-03 | 0 |
|  | *Moments* | **1.18e-01** | 2.16e-02 | **1.49e-03** | 0 |
| Selection, migration 3D,<br>T = 1.0 | $\partial a \partial i$ | 8.24e+01 | 4.19e-02 | 6.16e-04 | 0 |
|  | *Moments* | **5.02e+00** | **1.46e-02** | **1.73e-04** | 0 |
| Selection, migration 3D,<br>T = 5.0 | $\partial a \partial i$ | 4.00e+02 | 4.23e-02 | 3.35e-03 | 12 |
|  | *Moments* | **2.50e+01** | **1.36e-02** | **2.83e-04** | 0 |
| Selection, migration 3D,<br>T = 1.0 , neutral fs0 | $\partial a \partial i$ | 7.80e+01 | 2.98e-02 | 5.90e-04 | 3 |
|  | *Moments* | **5.01e+00** | **4.07e-03** | **8.63e-05** | 0 |
| Fast growth 3D,<br>T = 1.0 | $\partial a \partial i$ | 7.89e+01 | 3.48e+02 | 3.08e-01 | 7021 |
|  | *Moments* | **5.01e+00** | **7.07e-02** | **3.74e-03** | 0 |
| Out of Africa 3D | $\partial a \partial i$ | 5.65e+00 | 3.11e-02 | 2.52e-04 | 0 |
|  | *Moments* | **4.10e+00** | **1.57e-02** | **8.14e-05** | 0 |

**Table 7:** Performance comparisons between *∂a∂i* and *Moments* on several scenarios (80 samples per population). For *∂a∂i* simulations we used Richardson extrapolation to improve convergence. The time T provided is the simulation time in genetic units.

| Demographic model | method | exec time (s) | mean($\varepsilon_r$) | $\Delta$ LL | < 0 entries |
| --- | --- | --- | --- | --- | --- |
| Neutral equilibrium 1D | $\partial a \partial i$ | 6.91e-02 | 1.05e-02 | **5.44e-05** | 0 |
|  | *Moments* | **1.96e-03** | **1.01e-02** | 7.79e-05 | 0 |
| Neutral 1D,<br>T = 1.0 | $\partial a \partial i$ | 5.33e-02 | **4.07e-04** | **1.06e-07** | 0 |
|  | *Moments* | **1.42e-03** | 1.87e-02 | 5.77e-03 | 0 |
| Neutral 1D,<br>T = 5.0 | $\partial a \partial i$ | 1.61e-01 | **2.69e-04** | **2.46e-07** | 0 |
|  | *Moments* | **4.87e-03** | 4.63e-04 | 1.20e-05 | 0 |
| Selection 1D,<br>T = 1.0 | $\partial a \partial i$ | 5.25e-02 | **7.23e-04** | **4.88e-08** | 0 |
|  | *Moments* | **8.68e-03** | 1.87e-02 | 5.88e-03 | 0 |
| Selection 1D,<br>T = 5.0 | $\partial a \partial i$ | 1.70e-01 | 6.21e-04 | **9.79e-08** | 0 |
|  | *Moments* | **3.56e-02** | **5.09e-04** | 1.34e-05 | 0 |
| Neutral equilibrium 2D | $\partial a \partial i$ | 1.40e+00 | 1.04e-02 | 4.01e-03 | 15 |
|  | *Moments* | **8.86e-02** | **1.01e-02** | **1.56e-04** | 0 |
| Selection 2D,<br>T = 1.0 | $\partial a \partial i$ | 7.66e-01 | 1.04e-02 | **3.56e-03** | 0 |
|  | *Moments* | **2.48e-02** | **4.62e-04** | 1.01e-02 | 0 |
| Selection 2D,<br>T = 5.0 | $\partial a \partial i$ | 2.35e+00 | **4.53e-04** | 9.05e-03 | 0 |
|  | *Moments* | **1.07e-01** | 6.75e-04 | **-3.93e-03** | 0 |
| Selection, migration 2D,<br>T = 1.0 | $\partial a \partial i$ | 3.52e+00 | **1.53e-03** | **1.62e-06** | 0 |
|  | *Moments* | **2.43e+00** | 2.46e-03 | 5.53e-06 | 0 |
| Selection, migration 2D,<br>T = 5.0 | $\partial a \partial i$ | 1.61e+01 | 5.35e-03 | 6.25e-06 | 0 |
|  | *Moments* | **1.08e+01** | **4.68e-03** | **4.66e-06** | 0 |
| Fast growth 2D,<br>T = 1.0 | $\partial a \partial i$ | 3.56e+00 | 9.59e-02 | **2.59e-05** | 133 |
|  | *Moments* | **5.06e+00** | **6.13e-02** | 3.54e-05 | 0 |
| YRI-CEU 2D | $\partial a \partial i$ | **6.04e-01** | **1.06e-03** | **1.12e-06** | 0 |
|  | *Moments* | 1.63e+00 | 9.73e-03 | 2.14e-04 | 0 |
| Neutral equilibrium 3D | $\partial a \partial i$ | 1.95e+02 | 1.04e-02 | 1.19e-02 | 113549 |
|  | *Moments* | **4.35e+00** | **1.01e-02** | **2.34e-04** | 0 |
| Selection 3D,<br>T = 1.0 | $\partial a \partial i$ | 7.97e+01 | 9.15e-01 | 6.45e-03 | 0 |
|  | *Moments* | **9.63e-01** | **8.71e-06** | **-7.99e-03** | 0 |
| Selection 3D,<br>T = 5.0 | $\partial a \partial i$ | 2.82e+02 | 9.63e-01 | 1.69e-02 | 0 |
|  | *Moments* | **4.83e+00** | **1.25e-06** | **-6.29e-02** | 0 |
| Selection, migration 3D,<br>T = 1.0 | $\partial a \partial i$ | 8.37e+02 | 6.41e-03 | **9.26e-06** | 126 |
|  | *Moments* | **1.62e+02** | **4.46e-03** | 2.74e-04 | 0 |
| Fast growth 3D,<br>T = 1.0 | $\partial a \partial i$ | 8.31e+02 | 1.76e-01 | **5.01e-05** | 62559 |
|  | *Moments* | **1.59e+02** | **2.09e-02** | 2.50e-04 | 0 |
| Out of Africa 3D | $\partial a \partial i$ | **5.74e+01** | **5.90e-03** | **6.93e-06** | 102 |
|  | *Moments* | 1.06e+02 | 1.86e-02 | 1.29e-04 | 0 |

**Table 8:** Performance comparisons between *∂a∂i* and *Moments* on several scenarios (200 samples per population). For *∂a∂i* simulations we used Richardson extrapolation to improve convergence. The time T provided is the simulation time in genetic units.

| Demographic model | method | exec time (s) | mean($\varepsilon_r$) | $\Delta$ LL | < 0 entries |
| --- | --- | --- | --- | --- | --- |
| Neutral equilibrium 1D | $\partial a \partial i$ | 1.15e-01 | **1.03e-02** | **6.78e-05** | 0 |
|  | *Moments* | **3.20e-03** | 1.06e-02 | 9.62e-04 | 0 |
| Neutral 1D,<br>T = 1.0 | $\partial a \partial i$ | 1.16e-01 | **3.58e-05** | **4.41e-09** | 0 |
|  | *Moments* | **1.81e-03** | 2.04e-02 | 3.08e-02 | 0 |
| Neutral 1D,<br>T = 5.0 | $\partial a \partial i$ | 3.38e-01 | **2.35e-05** | **6.49e-09** | 0 |
|  | *Moments* | **7.83e-03** | 8.32e-04 | 6.73e-04 | 0 |
| Selection 1D,<br>T = 1.0 | $\partial a \partial i$ | 1.08e-01 | **6.45e-05** | **3.64e-09** | 0 |
|  | *Moments* | **1.50e-02** | 2.05e-02 | 3.17e-02 | 0 |
| Selection 1D,<br>T = 5.0 | $\partial a \partial i$ | 2.69e-01 | **5.53e-05** | **4.71e-09** | 0 |
|  | *Moments* | **4.02e-02** | 8.41e-04 | 7.41e-04 | 0 |
| Neutral equilibrium 2D | $\partial a \partial i$ | 8.37 | **1.02e-02** | 4.80e-03 | 13934 |
|  | *Moments* | **3.51e-01** | 1.06e-02 | **1.92e-03** | 0 |
| Selection 2D,<br>T = 1.0 | $\partial a \partial i$ | 4.69e+00 | 1.02e-02 | **4.06e-03** | 0 |
|  | *Moments* | **5.08e-02** | **2.04e-04** | 6.16e-02 | 0 |
| Selection 2D,<br>T = 5.0 | $\partial a \partial i$ | 1.29e+01 | **8.72e-04** | 8.91e-03 | 0 |
|  | *Moments* | **2.27e-01** | 1.77e-03 | **-2.42e-03** | 0 |
| Selection, migration 2D,<br>T = 1.0 | $\partial a \partial i$ | **1.84e+01** | **1.52e-03** | **4.71e-07** | 0 |
|  | *Moments* | 3.31e+01 | 3.67e-03 | 1.74e-05 | 0 |
| Selection, migration 2D,<br>T = 5.0 | $\partial a \partial i$ | **8.24e+01** | **8.23e-03** | **4.89e-06** | 18 |
|  | *Moments* | 1.58e+02 | 8.68e-03 | 1.14e-05 | 0 |
| Fast growth 2D,<br>T = 1.0 | $\partial a \partial i$ | **1.84e+01** | **1.56e-02** | **5.12e-06** | 1318 |
|  | *Moments* | 6.65e+02 | 6.11e-02 | 2.01e-05 | 0 |
| YRI-CEU 2D | $\partial a \partial i$ | **3.67e+00** | **3.41e-03** | **2.50e-06** | 0 |
|  | *Moments* | 1.54e+01 | 1.27e-02 | 2.53e-04 | 0 |
| Neutral equilibrium 3D | $\partial a \partial i$ | 3.71e+03 | **1.02e-02** | 1.42e-02 | 6325463 |
|  | *Moments* | **4.38e+01** | 1.06e-02 | **2.89e-03** | 0 |

**Table 9:** Test cases descriptions.

| Demographic model | Description |
| --- | --- |
| Neutral equilibrium kD | k constant size population(s), neutral simulation up to equilibrium (T=5.0) |
| Neutral kD | k growing population(s), neutral simulation. |
| Selection kD | k growing population(s) with selection: $Ns = 1$ and $h = 0.7$. |
| Selection kD, neutral fs0 | k growing population(s) starting at equilibrium with selection: $Ns = 1$ and $h = 0.7$. |
| Selection, migration kD | k growing populations with selection and migrations: $Ns = 1$, $h = 0.7$ and $m = 2.0$. |
| Selection, migration kD, neutral fs0 | k growing populations starting at equilibrium, with selection and migrations: $Ns = 1$, $h = 0.7$ and $m = 2.0$. |
| Fast growth kD | k exponentially growing populations with selection and migrations: $Ns = 1$, $h = 0.7$ and $m = 2.0$. |
| YRI-CEU | 2 populations Out of Africa expansion model, test case given in $\partial a \partial i$. |
| Out of Africa 3D | 3 populations Out of Africa expansion model as described in [9, 12] |

## F Additional features

*Moments* is based on *∂a∂i*’s interface but incorporates a new computation engine based on the method described in this paper. In addition, we added a few convenience features and optimizations. We developed a model plotting module that returns a schematic representation of a given demographic model. This is not only convenient for presenting results, but also proved very useful in troubleshooting. Figure 1 has been generated with this module. Moreover, it is now possible to directly import data and extract the allele frequency spectrum from a vcf file and to generate bootstrapped frequency spectra from the original data set. Finally, we implemented the Powell method for likelihood optimization, which we found to be more robust than the gradient and simplex methods used previously in *∂a∂i*, consistent with the ndings of [6].

